# Histopathological image QTL discovery of immune infiltration variants

**DOI:** 10.1101/126730

**Authors:** Joseph D. Barry, Maud Fagny, Joseph N. Paulson, Hugo J. W. L. Aerts, John Platig, John Quackenbush

## Abstract

Genotype-to-phenotype association studies typically use macroscopic physiological measurements or molecular readouts as quantitative traits. There are comparatively few suitable quantitative traits available between cell and tissue length scales, a limitation that hinders our ability to identify variants affecting phenotype at many clinically informative levels. Here we show that quantitative image features, automatically extracted from histopathological imaging data, can be used for image Quantitative Trait Loci (iQTL) mapping and variant discovery. Using thyroid pathology images, clinical metadata, and genomics data from the Genotype and Tissue Expression (GTEx) project, we establish and validate a quantitative imaging biomarker for immune cell infiltration. A total of 100,215 variants were selected for iQTL profiling, and tested for genotype-phenotype associations with our quantitative imaging biomarker. Significant associations were found in HDAC9 and TXNDC5. We validated the TXNDC5 association using GTEx *cis*-expression QTL data, and an independent hypothyroidism dataset from the Electronic Medical Records and Genomics network.

**One Sentence Summary:** We use a histopathological image QTL analysis to identify genomic variants associated with immune cell infiltration.

## Introduction

Genomics has had tremendous success in relating genetic variants to a variety of molecular phenotypes. Variants associated with gene expression, so-called expression Quantitative Trait Loci (eQTLs), are enriched for regions of active chromatin, influence gene regulation, and are involved in processes that contribute to disease [1, 2]. At the macroscopic level, genome-wide association studies (GWAS) that use physiological measurements of organ health have also provided insight into the genetic component of human disease. However, given the immense gap between SNP-level variation and tissue and body function, GWAS results concerning human traits can be challenging to interpret, in part due to the high correlations observed between studies of different traits [3]. This is especially true for autoimmune diseases, which have many shared genetic risk factors [4]. One way to increase GWAS variant interpretability and to discover new disease variants will be to place more focus on quantitative traits that lie at intermediate cellular and sub-tissue scales.

Histopathology has for decades remained the standard for the diagnosis and grading of many complex diseases. Advances in digital slide imaging have allowed for unprecedented resolution at tissue, cellular, and sub-cellular scales. As such, histopathology could provide the genomics community with a vast resource of quantitative traits for evaluating disease phenotype across a range of mesoscopic scales. Previous histopathological GWA studies have used discrete pathology grading schemes as quantitative traits. While such schemes are highly effective for guiding clinical decisions on the level of individual patients, and can in principle identify disease variants in GWAS [5], they do not scale well to the hundreds or thousands of samples needed due to inter- and intra-observation bias [6-8]. Unbiased, automatically extracted, continuous features have been shown to improve the predictive power of survival analyses with pathology data in comparison to discrete grades [9, 10]. GWA studies using quantitative image features extracted from radiological imaging data have successfully identified COPD-relevant variants using Computed Tomography [11], and variants related to Alzheimer’s Disease and Mild Cognitive Impairment using Magnetic Resonance Imaging [12]. Thus, genome association studies that leverage automated imaging analysis methodologies can be highly effective. With quantitative image analysis now routinely applied to digital pathology datasets [9, 10, 13-15], it is clear that unbiased, continuous image features can readily be extracted for histopathological GWAS variant discovery.

Nevertheless, little attention has been paid to the use of cellular imaging to obtain quantitative traits for understanding the connection between genotype and cellular phenotypes. Part of the reason for this may be that there are few available datasets for which both genomic profiling and standardized histopathology data are available for any appreciable number of samples. The Genotype-Tissue Expression (GTEx) resource [16, 17] has collected germline genotype data from over 450 autopsy research subjects, as well as standardized histological imaging and RNA-Seq gene expression from approximately 50 different body sites.

One of the tissues for which there is substantial histological imaging data is the thyroid. The thyroid plays a central role in the endocrine system, producing thyroid hormones that influence protein synthesis and metabolism. Hashimoto Thyroiditis (HT) is an autoimmune disease affecting approximately 5% of the population, in which the thyroid is slowly destroyed, frequently leading to hypothyroidism, particularly in women [18]. Among the GTEx population there are 341 individuals for whom paired thyroid imaging and gene expression data was available; of these 31 had morphological evidence of HT according to GTEx pathology notes. Leukocyte infiltration is a morphological hallmark of HT, and necessary for disease progression. We saw this dataset as an ideal place to test the hypothesis that cellular imaging features could be used to discover immune cell infiltration variants in an image Quantitative Trait Locus (iQTL) analysis.

### Quantitative image analysis

Image data was downloaded from the GTEx histological image archive, and convolved with a Gaussian filter to smooth out pixel-level variation on a length scale smaller than observed regions of leukocyte invasion in HT samples. Color deconvolution to estimate the Hematoxylin and Eosin channels directly was not performed to avoid the risk of introducing any biases from the deconvolution process that might negatively affect downstream QTL fits. Individual tissue pieces were segmented using adaptive thresholding (Figure 1A), and 117 Haralick image features were extracted for each tissue piece using the Bioconductor image processing package *EBImage* [19] (Transparent Methods). To capture patterns across multiple length scales, we incorporated three Haralick scales that sampled every 1, 10, or 100 pixels. Haralick features are widely used for the quantification of image texture, and take into account correlations between neighboring pixels in an image [20]. Given that lymphocyte invasion has a profound effect on histological image texture, we reasoned that Haralick features could be ideal candidates for capturing HT cellular phenotype. After removing overly small tissue pieces, averaged image feature values were calculated across pieces to give a single value for each sample and feature. In preparation for downstream model fitting, features were log_2_ transformed, and centered and rescaled using a Z-score to ensure feature comparability across samples (Figure 1B). The mean and standard deviation values used for the Z-score were saved for use in later analyses. A fully worked through image analysis example with the applied segmentation parameters is supplied as part of a supplemental R data package.

**Figure 1.**
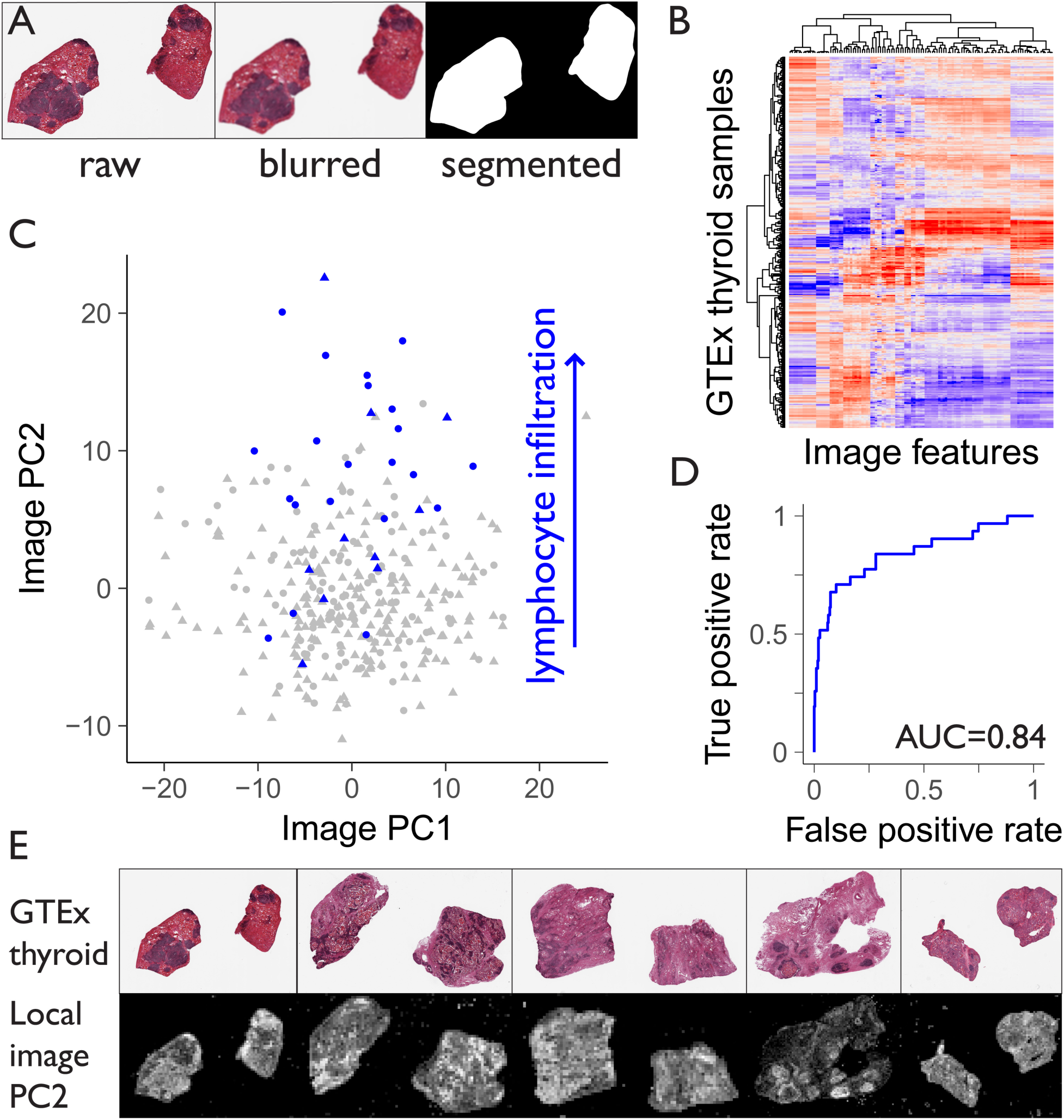
Thyroid image processing and establishment of a quantitative imaging biomarker for immune cell infiltration. (**A**) Digital pathology slide for thyroid sample GTEX-11NV4. Raw: pre image processing. Blurred: post Gaussian convolution. Segmented: tissue piece masks after adaptive thresholding. (**B**) Heatmap of 117 log_2_ transformed and standardized Haralick image features against 341 thyroid samples. (**C**) PC1 versus PC2 from a PCA of the image feature matrix. Blue points indicate patients with Hashimoto Thyroiditis, as identified from pathology notes. Circles indicate females, and triangles males. (**D**) ROC showing biomarker performance of PC2 for predicting HT. (**E**) Top row, five thyroid samples with highest values of image PC2. Bottom row, local image PC2 signal. Bright regions correspond to high local image PC2. Image brightness has been rescaled to aid visualization.

To reduce the number of candidate features and to identify the primary directions of variability in our image feature matrix, we performed Principal Component Analysis (PCA), and found that the first two imaging Principal Components (PCs) were sufficient to explain 73% of the variance. By inspection of the images, low values of PC1 were found to be associated with interior holes in the thyroid pieces (Figure S1A), an effect we adjudged to be primarily technical in origin. High values of image PC2 were visually strongly associated with the presence of invading leukocytes (Figure S1B).

For validation of image PC2 using clinical metadata, we identified all samples for which a HT phenotype was indicated in the comments of the original GTEx pathologists (Figure 1C blue points). We note that the morphological phenotype of HT is not definitive evidence of clinical HT, which requires assessment of thyroid hormone levels, ultrasonographic scanning, and additional clinical findings [21]. A ROC analysis showed that image PC2 was a highly performant biomarker for HT phenotype, with an AUC of 0.84 (Figure 1D). Using logistic regression, we found that HT phenotype was significantly associated with image PC2 (OR=1.3, P=1.4×10^−9^) after correcting for sex and age (Transparent Methods). A higher association with HT phenotype for women was also observed after correcting for age (OR=4.2, P=4×10^−4^), a known clinical characteristic of the disease [18]. That image PC2 is a performant biomarker for HT phenotype likely does not have direct clinical application since a pathologist can easily identify HT morphology by inspection of pathology images; we considered its utility to be for downstream integration with genomic data.

Since the fibrous variant of clinical HT accounts for up to 10% of cases [22] we investigated co-occurrence of HT and fibrotic phenotypes in GTEx pathology notes. We found that 9/31 samples with HT phenotypes also had fibrotic phenotypes, which was significantly enriched compared to the background set of non-HT samples (Fisher Test; P=0.002, OR=4.2). However, samples with both HT and fibrotic phenotypes did not show any clear classification pattern in the image PC1 vs PC2 plot (Figure S2A) that would suggest segregation by disease subtype.

Next we inspected PC2 feature loadings from the image PCA (Table 1). These represented the weighted contribution of each Haralick feature to image PC2. We found that no single feature had a notably higher magnitude of loading than any other, showing that signals from many different Haralick features were required (Figure S2B). To assess the relative contribution of each RGB color channel we summed the absolute value of PC2 principal component loadings for all features extracted from each channel. Proportionately, we found that the red channel features contributed 47%, the green channel 28%, and the blue channel 26%, to the image PC2 signal. This was suggestive of Eosin contributing the most to the observed signal.

**Table 1.** Principal component loadings for image PC2 are supplied in the supplementary data table pc2loading.tsv. Labeling follows the nomenclature [*channel*].h.[*feature*].s[*scale*] where *channel* indicates the RGB color channel, *feature* is one of Haralick’s 13 textural features, and *scale* gives the length scale over which features are assessed (1, 10, or 100 pixels; Transparent Methods).

To verify that image PC2 signal was originating from regions of immune infiltrate, we mapped image PC2 signal to tiles of size 250×250 pixels using the following procedure. Haralick features were extracted within each tile, log_2_ transformed, and centered and rescaled using a Z-score. Crucially, the mean and standard deviation values used for the Z-score were those calculated previously when estimating feature variability across samples. The resulting feature values were then multiplied by their corresponding image PC2 component loading (Figure S2B), and summed together to give the final estimate for local image PC2. Figure 1E shows local image PC2 signal for the five samples with highest image PC2 signal in Figure 1C. Bright regions indicate where local image PC2 signal is highest (Figure 1E bottom row), and can be seen to correspond directly to dark regions of dense immune infiltrate in the original images (Figure 1E top row).

### Integrated RNA-Seq and image analysis: imaging biomarker validation

Next we analyzed GTEx gene expression data associated with the 341 thyroid images. After the removal of lowly expressed genes, counts were log_2_ transformed and underwent quantile normalization to ensure comparability of samples (Transparent Methods). To test for associations between gene expression and image PCs, linear models were used to fit each image PC against normalized gene readouts while correcting for age, sex, sample collection site, tissue autolysis score, and RNA extraction type. To remove bias caused by unknown confounders (Figure S3A) we fitted 20 PEER (Probabilistic Estimation of Expression Residuals) factors [23] to the gene expression data while including the known confounders in the PEER fit; a similar approach was used by the GTEx consortium [16, 24]. The first PEER factor was not included as a covariate in the linear model fits due to the strength of its correlation with our image PC2 biomarker (Pearson r=0.53, P<2.2×10^−16^), which indicated that it likely contained valuable biological information. While other PEER factors may capture both biological signal and systematic noise, their inclusion as covariates successfully removed the technical bias observed in Figure S2A. The results from fits correcting for the known confounders as well as the 19 remaining PEER factors revealed that image PC2 was systematically and highly associated with gene expression (Figure 2A), suggesting that leukocyte infiltration is at least a major, if not the primary source of expression variability in GTEx thyroid tissue. In contrast, image PC1 had comparatively few significant associations with thyroid gene expression (Figure 2A).

**Figure 2.**
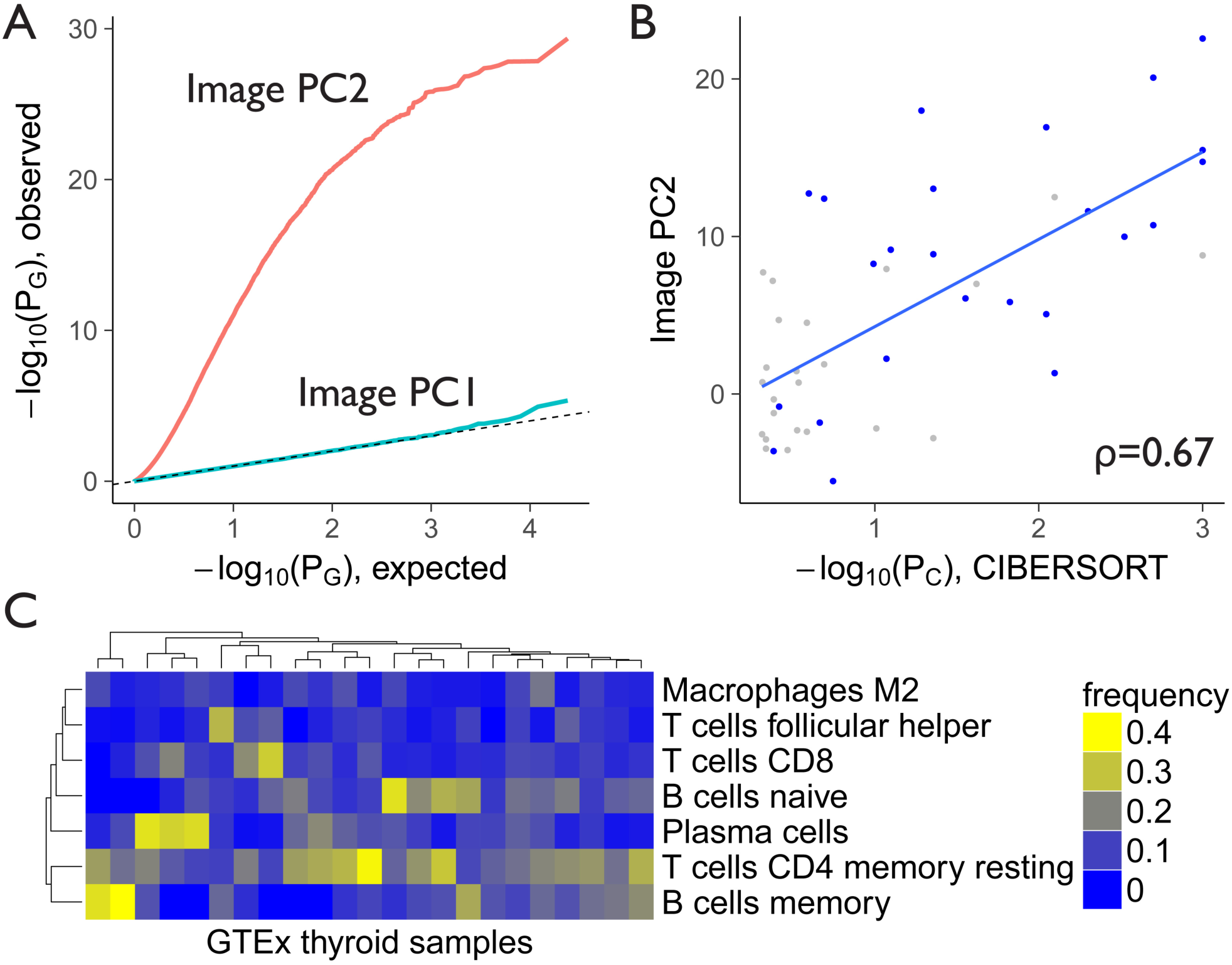
Integration of imaging data with gene expression analyses verifies that image PC2 is highly associated with immune cell infiltration. (**A**) A QQ plot of p-values (P_G_) from the regression analysis of image PC1 and PC2 against thyroid gene expression for 23,993 genes. (**B**) Correlation of image PC2 with –log_10_(P_C_) from the CIBERSORT analysis for samples with P_C_<0.5. Blue points are samples for which Hashimito Thyroiditis was indicated in GTEx pathology notes. (**C**) Frequencies of immune cell types reported from CIBERSORT for samples with P_C_<0.1. Cell types with an average frequency of 5% or more are shown.

To identify which biological pathways may drive the significant image PC2 associations with gene expression we performed gene-set enrichment analysis, choosing as a test set 2,913 genes with –log_10_(P_G_) >10 in the gene expression analysis and all other 21,080 genes as a background set (Transparent Methods). Significant enrichment was observed for a number of gene ontology terms related to immune function, and the activation and signaling activity of invading T and B lymphocytes (Table 2).

**Table 2.** Significant gene ontology terms for genes highly associated with image PC2 (−log_10_(P_G_)>10, see Figure 2A) in the gene expression regression analysis. All terms with an odds ratio of greater than 5 and an adjusted p-value of less than 0.01 are shown. BP: Biological Process, CC: Cellular Component.

| GO id | Type | Description | Odds | P | P<sub>adj</sub> | N |
| --- | --- | --- | --- | --- | --- | --- |
| GO:0050776 | BP | regulation of immune response | 6.10 | 1.3E-14 | 5.8E-11 | 43 |
| GO:0031295 | BP | T cell costimulation | 5.83 | 6.0E-11 | 2.1E-07 | 32 |
| GO:0042110 | BP | T cell activation | 5.57 | 2.6E-08 | 3.6E-05 | 24 |
| GO:0042113 | BP | B cell activation | 8.67 | 7.0E-08 | 8.6E-05 | 17 |
| GO:0045060 | BP | negative thymic T cell selection | 25.51 | 6.7E-07 | 5.4E-04 | 10 |
| GO:0050690 | BP | regulation of defense response to virus by virus | 6.96 | 2.1E-06 | 1.5E-03 | 15 |
| GO:0042101 | CC | T cell receptor complex | 17.00 | 2.5E-06 | 1.6E-03 | 10 |
| GO:0050853 | BP | B cell receptor signaling pathway | 5.89 | 7.2E-06 | 3.8E-03 | 15 |

To confirm the association between image PC2 and the presence of invading immune cells on the level of individual samples, we ran our gene expression data through CIBERSORT, an algorithm designed to deconvolve complex cell mixtures [25]. The LM22 leukocyte gene signature matrix supplied by the CIBERSORT team was used to perform the deconvolution as it could in principle detect and distinguish between a number of T and B cell types. As expected, CIBERSORT failed to detect immune cells in the majority of healthy thyroid samples. However, in samples where immune cells were detected (CIBERSORT P_C_<0.5), the significance of detection correlated significantly with image PC2 (Figure 2B; Spearman ρ=0.67, P=3×10^−7^). Figure 2C shows the cell type signature for samples with P_C_<0.1 and cell types with an average CIBERSORT frequency of more than 5% across samples. This analysis confirmed the presence of T-cell CD4 memory resting, T-cell CD8, naïve and memory B cells, as well as other immune cell types, on the level of individual samples.

The CIBERSORT algorithm also detected infiltrating immune cells in several samples for which image PC2 was high but HT phenotype was not indicated by GTEx pathologists (Figure 2B grey points), suggesting that our imaging biomarker may be able to detect a broader spectrum of thyroid autoimmune processes. By inspection, we confirmed that many of these images did show infiltrative phenotypes but did not exhibit the classical HT visual phenotype (for example GTEX-OXRO, GTEX-144FL). One possible clinical application of our imaging approach would therefore be to automatically flag such samples for rechecking by pathologists.

To demonstrate the advantage of using PCA for the identification of a candidate imaging biomarker, as opposed to directly using the individual Haralick image features, we correlated each of the 117 image features with CIBERSORT significance of detection for immune cells, in the same manner as above. Image PC2 had a higher Spearman correlation coefficient than all image features, with the closest having a coefficient of ρ=0.60 (Figure S2B). Thus, image PC2 combined information from multiple image features to attain a higher overall performance.

### iQTL identification of variants associated with immune infiltration

Based on our analysis of image features and gene expression, we decided to use image PC2 as a quantitative trait capturing the cellular features of immune cell infiltration in an image QTL analysis. Since we did not have sufficient sample size to achieve high power in a genome-wide iQTL analysis that would include millions of variants, and other suitably large datasets were not available, we decided to reduce the search space by asking a more targeted question of the data.

Since leukocyte invasion was associated with a higher prevalence of differentially upregulated genes (Figure 3A) we hypothesized that the genotype of upregulated genes in GTEx samples with a HT phenotype could in part determine the extent of leukocyte infiltration. Therefore, the analysis was further restricted to 100,215 candidate SNPs residing in 1,380 genes that were significantly (-log_10_(P_adj_) >7, Figure S4A) and positively (log_2_(Fold-Change) >0.5) differentially expressed in 31 HT cases versus 310 controls (Figure 3A blue points) using *DESeq2* [26]. We note that HT phenotype, as indicated by GTEx pathologists, was not included in the feature set used to build our imaging biomarker. Therefore, the SNP selection procedure was independent of the biomarker subsequently used as the quantitative trait in the iQTL analysis.

**Figure 3.**
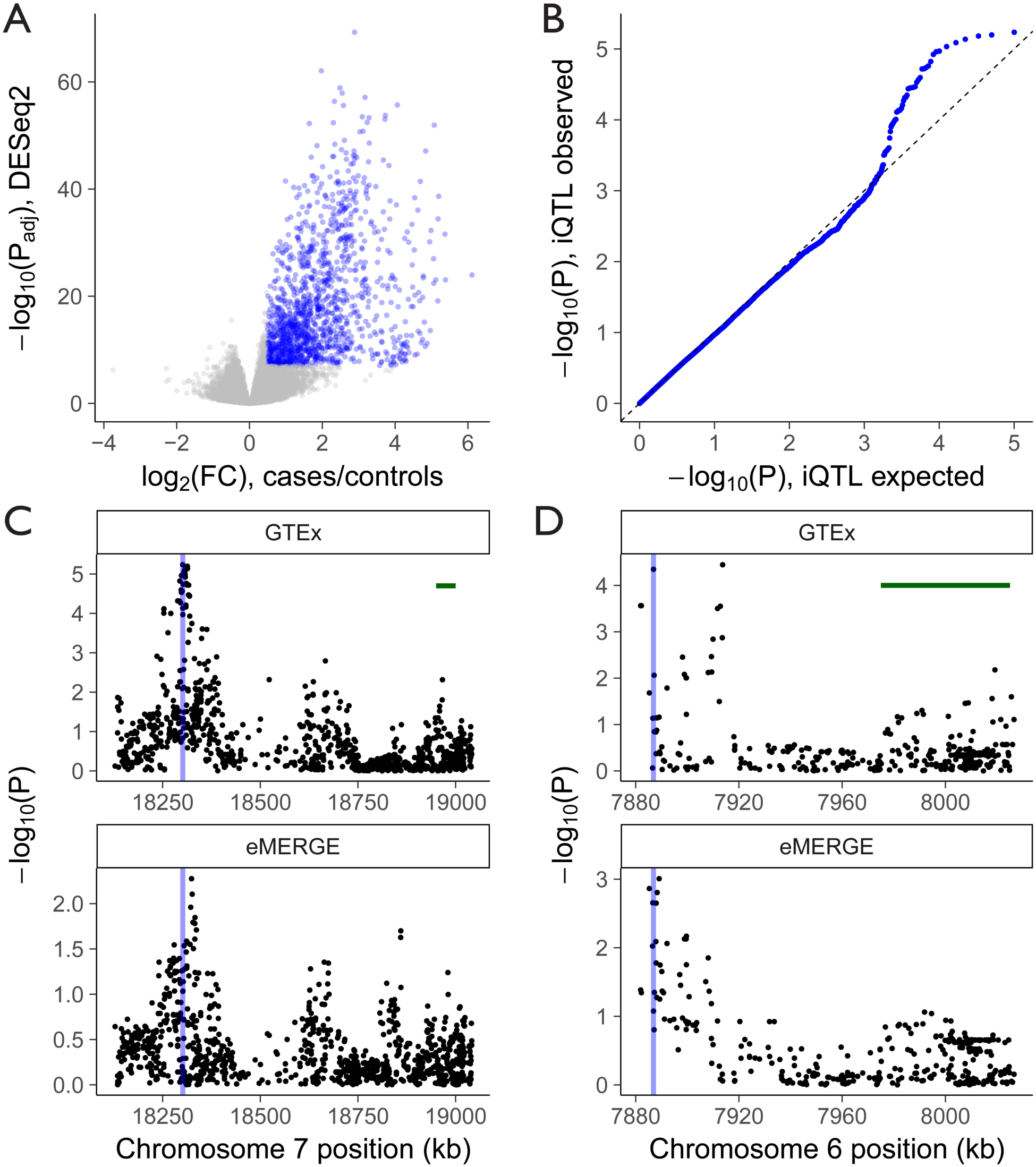
An image QTL analysis finds associations between genomic variants and our image PC2 biomarker for thyroid autoimmune disease. (**A**) Selection of 1,380 candidate genes (blue points) based on their positive fold-change (log_2_(FC)>0.5) and significant differential expression (-log_10_(P_adj_)>7) in GTEx samples with Hashimoto Thyroiditis (HT) phenotypes. (**B**) A QQ plot showing expected vs observed p-values from image QTL fits of 100,215 candidate SNPs residing in the selected genes highlighted blue in panel A. (**C-D**) –log_10_(P) vs genomic coordinates for GTEx iQTLs (top panels) and eMERGE variant association with HT (bottom panels) for all tested SNPs in HDAC9 (C) and TXNDC5 (D). Blue vertical lines indicate the locations of the most significant SNP for each gene after multiple testing correction using the IHW method described in the main text. Plot ranges are mapped to the start and end positions of the genes, as defined by GTEx-consortium transcript data. Horizontal green bars are of length 50kb.

We then limited our analysis to the 292 samples that had both thyroid imaging and genotype information. The genotype data had already undergone rigorous quality control and filtering by the GTEx consortium, including but not limited to Hardy-Weinberg equilibrium and imputation quality criteria. Due to sample size considerations, minor allele frequencies (MAFs) of less than 10% were removed, as well as SNPs with a missingness frequency of more than 10%.

Image QTL linear model fits were implemented using the R package *MatrixEQTL* [27] by treating image PC2 as a pseudo *trans* gene. Fits were corrected for age, sex, race, ancestry, processing center, and tissue autolysis score (Transparent Methods). A QQ plot comparing the distribution of observed p-values to the expected uniform distribution showed that the data was well behaved under the null hypothesis (Figure 3B), with a departure from the dashed line for a group of SNPs with low p-values. Independent Hypothesis Weighting (IHW) [28] using MAF as the independent covariate was used to correct for multiple testing. In a data-driven way, this method assigned different weights to variants based on their MAF (Figure S4B) to maximize the number of null hypothesis rejections while controlling for Type I error. A total of 21 SNPs across 3 haplotype blocks were identified as being significant (P_IHW_<0.05; Table S1). As expected, IHW attained higher power as compared to standard, unweighted Benjamini and Hochberg FDR (Table S1).

Strikingly, 20 of the 21 significant SNPs resided in Histone Deacetylase 9 (HDAC9), an enzyme linked to epigenetic control of gene transcription, and previously proposed to be an epigenetic switch for T-cell mediated autoimmunity [29]. Histone deacetylase inhibitors have been effective in the treatment of hypothyroidism in mice [30], and autoimmune diseases in general [31]. A plot of SNP significance along HDAC9 revealed a sharp peak at 18,301 kb on chromosome 7 (Figure 3C). To our knowledge this is the first reported association between HDAC9 variants and thyroid immune cell infiltration. A search of GWAS Central (www.gwascentral.org [32]) identified modest associations between Ulcerative Colitis (UC) and two of our significant image QTL variants (-log_10_(P)=2.3 for *rs215122*, -log_10_(P)=2.7 for *rs2529749*) [33]. While further work will be required to investigate the biological effects of HDAC9 genotype in both UC and HT, a common genetic basis would be consistent with evidence that these conditions present concurrently at higher rates than would be expected in the general population [34].

One variant in the Thioredoxin Domain Containing 5 gene (TXNDC5), *rs11962800*, was found to be significant at the level of P_IHW_<0.05 (Table S1). TXNDC5 expression is induced by hypoxia [35], and variants in this gene have been associated with a number of autoimmune diseases including rheumatoid arthritis [36] and vitiligo [37]. As for HDAC9, variants in TXNDC5 have not previously been associated with immune cell infiltration, again suggesting that iQTLs can uncover clinically relevant associations. It should be noted that *rs11962800* was reported as a *cis*-expression QTL in thyroid tissue by the GTEx consortium (P=7.7×10^−21^; www.gtexportal.org [24]). Since our analysis shows that TXNDC5 expression is associated with leukocyte invasion in thyroid tissue, it is possible that any expression-based result could be biased by cell-type mixing, as has been highlighted in GTEx lung tissue [38]. However, *rs11962800* was also a reported eQTL in 16 other GTEx tissues [24], with thyroid ranked 3/17 in terms of highest significance. Next we relaxed the IHW adjusted p-value cutoff for iQTLs to 0.4, and found that the top five iQTLs in TXNDC5 were also eQTLs (Table S2). Additionally, 12/31 of the GTEx TXNDC5 eQTLs overlapping with our set of tested variants were iQTLs or in linkage disequilibrium with iQTLs (Figure S5 red points). Together, these results offer the mechanistic hypothesis that genetic variation in TXNDC5 affects HT imaging phenotype directly through *cis*-regulation of RNA expression levels.

Next we asked how the performance of our imaging biomarker might compare with expression-based biomarkers for immune infiltration variant discovery. We chose the first expression PEER factor (PEER1) as a comparative candidate biomarker based on the earlier observation that it was significantly correlated with image PC2. After verifying that PEER1 was performant for predicting HT phenotype (AUC 0.88, Figure S6A), we ran another QTL analysis using the same 100,215 variants as for the iQTL analysis, but with PEER1 as the response variable. A QQ plot revealed that PEER1 performed poorly (Figure S6B), with no significant QTL hits. Therefore, image PC2 was in this case a superior choice of biomarker for variant discovery.

### Validation of iQTL results using an independent dataset

To attempt validation of the observed associations between HDAC9 and TXNDC5 genotype, and thyroid immune cell infiltration in an independent dataset, we obtained genotype and phenotype data, and clinical metadata from the Electronic Medical Records and Genomics (eMERGE) network [39] (Transparent Methods). This dataset has been used previously for hypothyroidism variant discovery [40]. Criteria for defining HT cases included the assessment of TSH and FT4 levels (Transparent Methods). While levels of biochemical markers are not necessarily correlated with morphological grading of HT, immune cell infiltration is pathognomonic of the disease [21]. An increased prevalence of immune cell infiltration is therefore likely to be present in a large set of HT cases vs controls.

We selected 1,261 chronic autoimmune hypothyroidism (presumptive HT) cases and 4,457 non-hypothyroidism controls (Transparent Methods). As over 96% of the hypothyroidism cases were from the Caucasian cohort, and 250/292 of the GTEx samples used for the image QTL analysis were of European American descent, to simplify the analysis all non-Caucasian cohorts were removed, leaving 1,213 cases and 3,789 controls. For consistency with the image QTL analysis, SNPs with a minor allele frequency of less than 10% and a missingness frequency of more than 10% were removed. One SNP in HDAC9 was removed due to deviation from Hardy-Weinberg equilibrium (Transparent Methods).

We used logistic regression to test for associations between hypothyroidism status, and HDAC9 and TXNDC5 genotype while correcting for patient age (decade of birth), sex, collection site, and ancestry (Transparent Methods). For both genes, the location of the most significant GTEx image QTL peaks, as determined by their IHW-adjusted p-values, coincided closely with the highest eMERGE peaks (Figure 3C,D vertical lines). While a colocalisation analysis did not identify a single causative SNP common to both datasets, we noted a high degree of similarity between the association profiles. To quantitatively assess the concordance of the association signatures between the GTEx and eMERGE data, we first selected the SNPs common to both datasets, and grouped neighboring SNPs into bins of size 20 SNPs based on their genomic coordinates. The lowest p-value in each bin was then selected for both datasets. A correlation test revealed that the association signatures between eMERGE and GTEx data were significantly concordant across both HDAC9 (Figure S7A1; Spearman ρ=0.59, P<2.9×10^−6^) and TXNDC5 (Figure S7A2; Spearman ρ=0.73, P=9×10^−4^).

Next we checked if linkage disequilibrium (LD) could be artificially driving the observed concordance. SNPs were pruned for LD using PLINK (Transparent Methods), and the binning and correlation analysis was repeated. To account for the reduction in the number of SNPs (HDAC9, 1071 to 330; TXNDC5, 347 to 88), smaller bin sizes of 6 and 5 SNPs were used for HDAC9 and TXNDC5 respectively. A correlation test revealed a more modest but still significant correlation for HDAC9 (Spearman ρ=0.30, P=0.026), and a significant correlation for TXNDC5 (Spearman ρ=0.72, P=0.001). Thus, HDAC9 concordance was partially biased by LD, while TXNDC5 concordance was unaffected.

To verify that it was hypothyroidism status driving the concordance, we performed 1,000 permutations of the hypothyroidism case/control labels in the eMERGE dataset and repeated the logistic regression, binning, and correlation analyses between the LD-pruned GTEx and eMERGE data. The resulting permutation p-value did not reach significance for HDAC9 (P=0.127), likely due to the lower nominal p-values remaining after the LD-pruning. However the TXNDC5 permutation value was significant (P=0.006), demonstrated that it was unlikely to obtain the observed TXNDC5 concordance between eMERGE and GTEx profiles by chance (Figure S7B1,2).

While we validated our TXNDC5 association using eMERGE hypothyroidism status, there are limitations to the validation approach. First, the analysis identified common regions of association, rather than individual causative SNPs. The colocalisation analysis was likely hindered by the modest GTEx sample size, and future iQTL studies may require higher numbers of samples to identify causative SNPs. Second, the severity of the HT morphological phenotype and clinical hypothyroidism status are not necessarily correlated [21], although immune cell infiltration is a prerequisite for clinical HT. This may in part explain why higher nominal p-values were observed in the GTEx data. Another explanation might be that quantitative histological image analysis captures subclinical, as well as clinical disease phenotype as a smooth function, while binary categorizations of disease status do not take into account informative subclinical disease cases, or the morphological severity of the disease phenotype.

### Outlook

Our intuition, along with over a decade of progress in GWAS, tells us that genotype and phenotype are associated in ways that can be quantitatively assessed. The growth of integrated data resources such as GTEx, and the advancement of digital slide imaging technologies, provides the opportunity to explore that association at the cellular level. This study demonstrates the potential of histopathological image-based QTL profiling for *de novo* discovery of disease variants, especially when unbiased, continuous, automatically extracted image features are used as quantitative traits. Our results indicate that iQTLs can identify different variants to cases vs controls GWAS, offering a complementary approach that leverages cell-level phenotypes for target discovery. While we applied iQTLs to a thyroid dataset, the method is completely generalizable to other tissues and disease contexts. With larger sample sizes and study designs tailored for specific diseases, iQTL scans can be extended to millions of variants. Ultimately, digital pathology image analysis approaches could revolutionize genome-wide association studies by providing a wealth of unbiased quantitative traits at the cellular level that have high levels of biological interpretability.

## Notes

Code and analysis reproducing paper figures, as well as reported image and statistical analyses performed on non-protected data, are supplied as part of an R package hosted on github: https://github.com/josephbarry/iQTL.

## Author Contributions

JDB, HJWLA and JQ conceived of the project. JDB performed all image and data analysis, with input from MF (CIBERSORT analysis), JNP (statistical analysis), and JP (QTL pipeline). JDB wrote the paper with input from all authors.

## Acknowledgements

This work was supported by grants from the US National Institutes of Health, including 5U01CA190234 and 5U24CA194354 (JDB, HJWLA, JQ), 5R01AI099204 (JP, JQ); 5R01HL111759 (JQ, JNP), and 5P50CA127003 (JQ). This work was conducted under dbGaP approved protocols #9112 and #13896.

---

**Figure S1.**
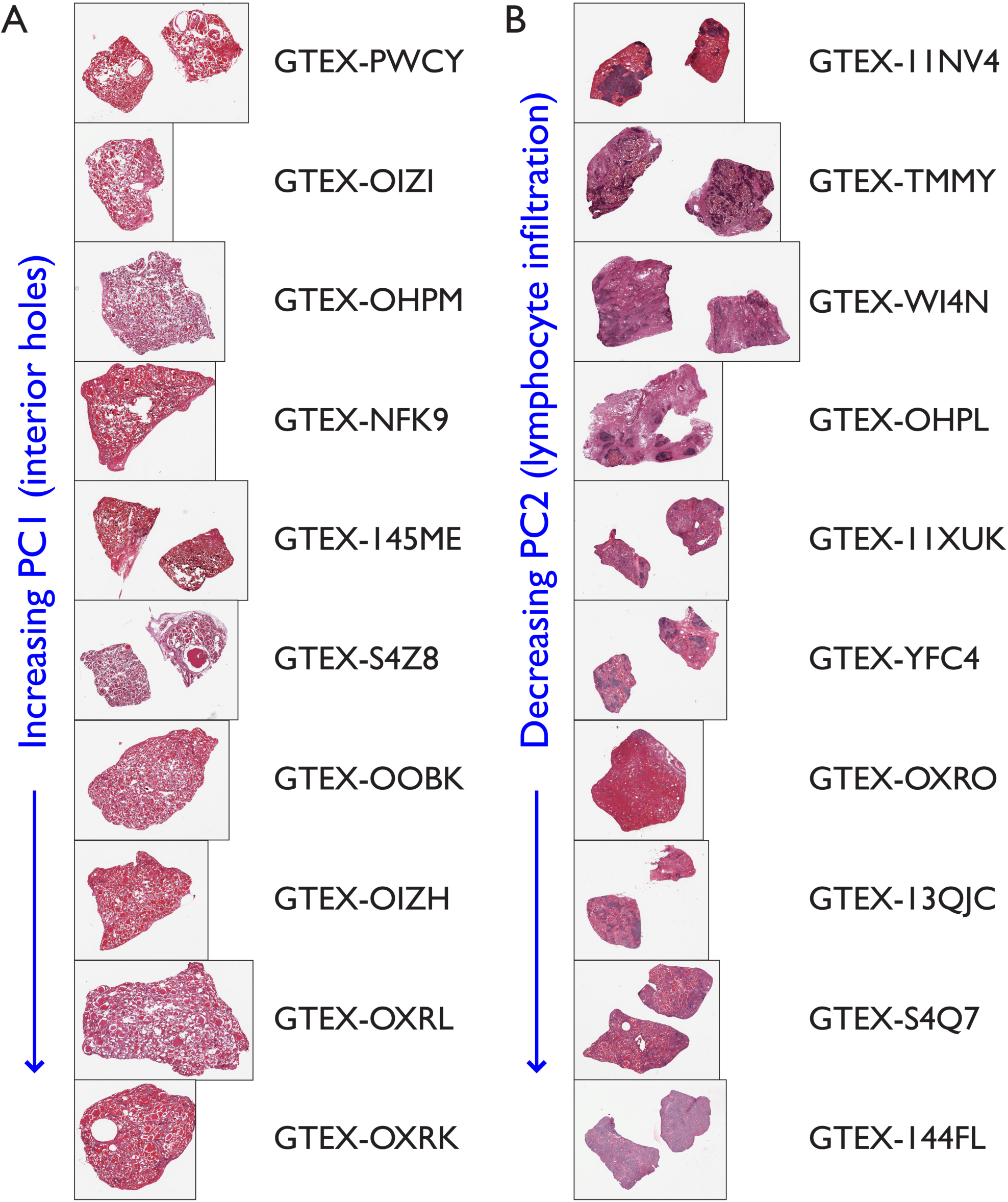
Examples of phenotypes associated with image PC1 and PC2. (**A**) Images with the 10 lowest values of PC1. Interior holes were observed, many of which were likely associated with tissue damage incurred during sample preparation. (**B**) Images with the 10 highest values of PC2. Lymphocyte invasion phenotypes were apparent.

**Figure S2.**
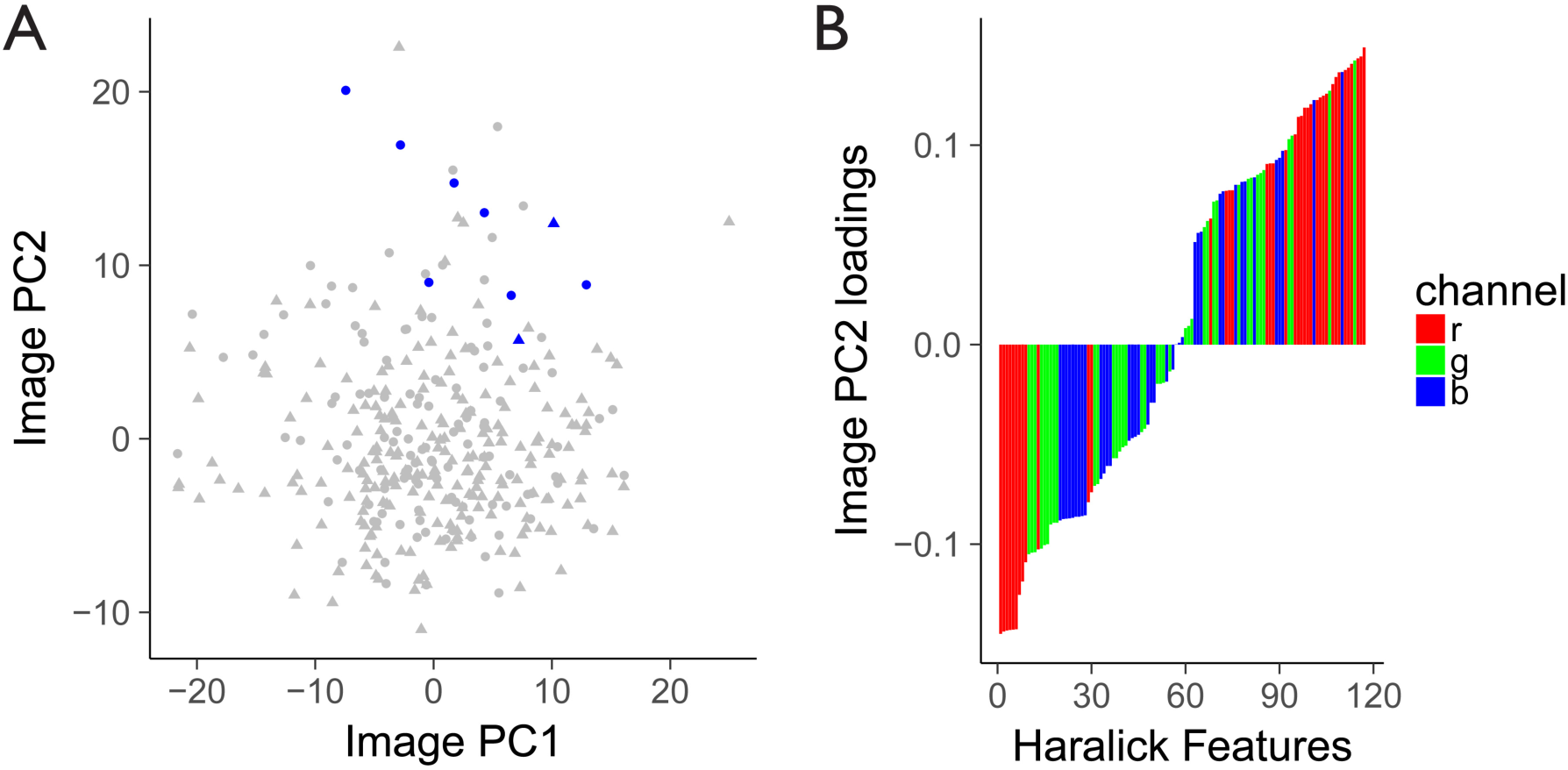
(**A**) PC1 versus PC2 from a PCA of the image feature matrix. Blue points indicate patients with both Hashimoto Thyroiditis and fibrotic phenotypes, as identified from pathology notes. Circles indicate females, and triangles males. (**B**) Haralick feature loadings from the second principal component of the image PCA, ranked by increasing value. Colors indicate RGB channels.

**Figure S3.**
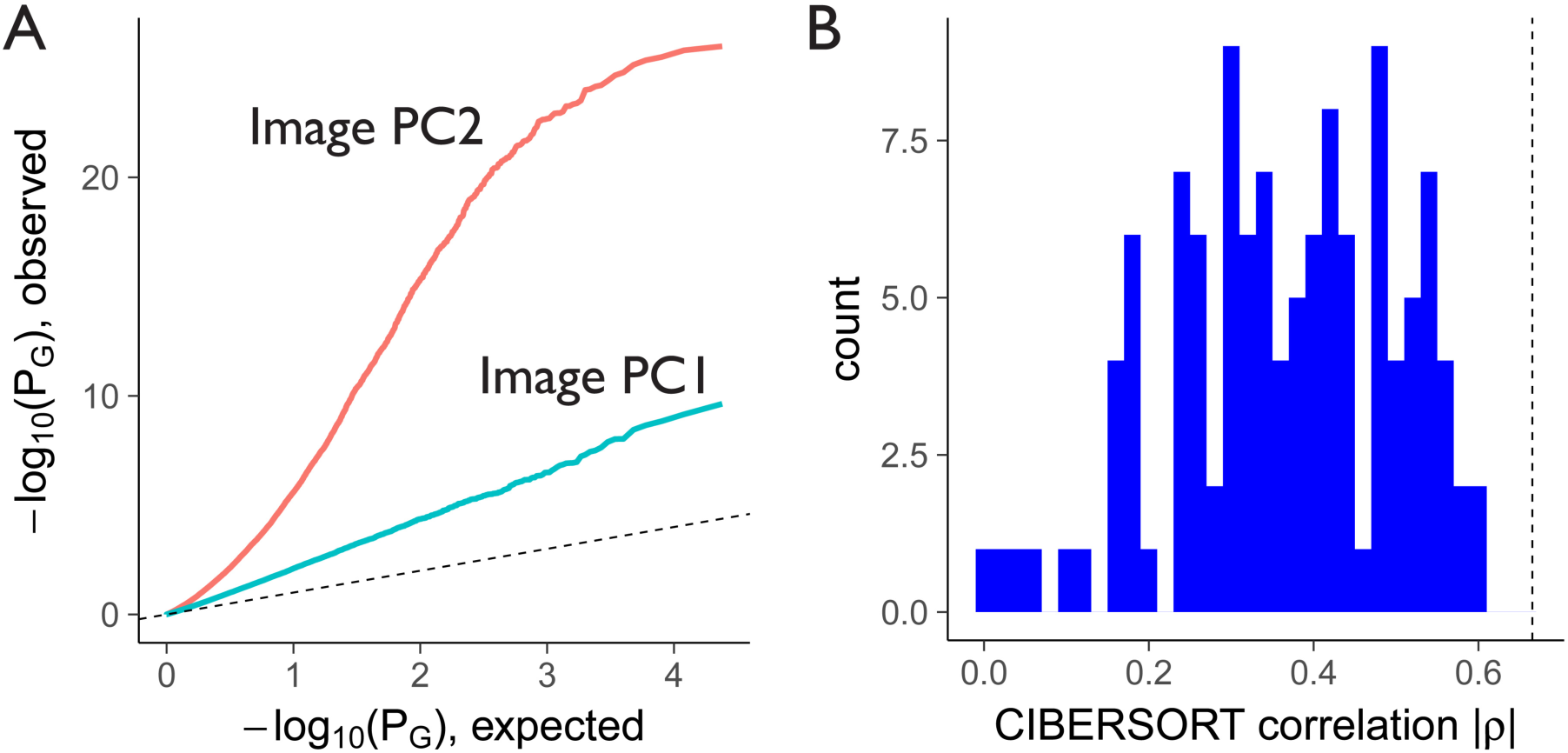
(**A**) QQ plot showing deviation from the dashed line (of slope 1) when fitting image PCs against gene expression while only correcting for the known confounders listed in the main text. Additionally including PEER factors as covariates removed the observed bias (Figure 2A). (**B**) Histogram of the absolute value of Spearman correlation coefficients between all 117 Haralick image features, and CIBERSORT significance of detection for immune cells. Dashed line indicates the Spearman correlation coefficient for image PC2 (ρ=0.67).

**Figure S4.**
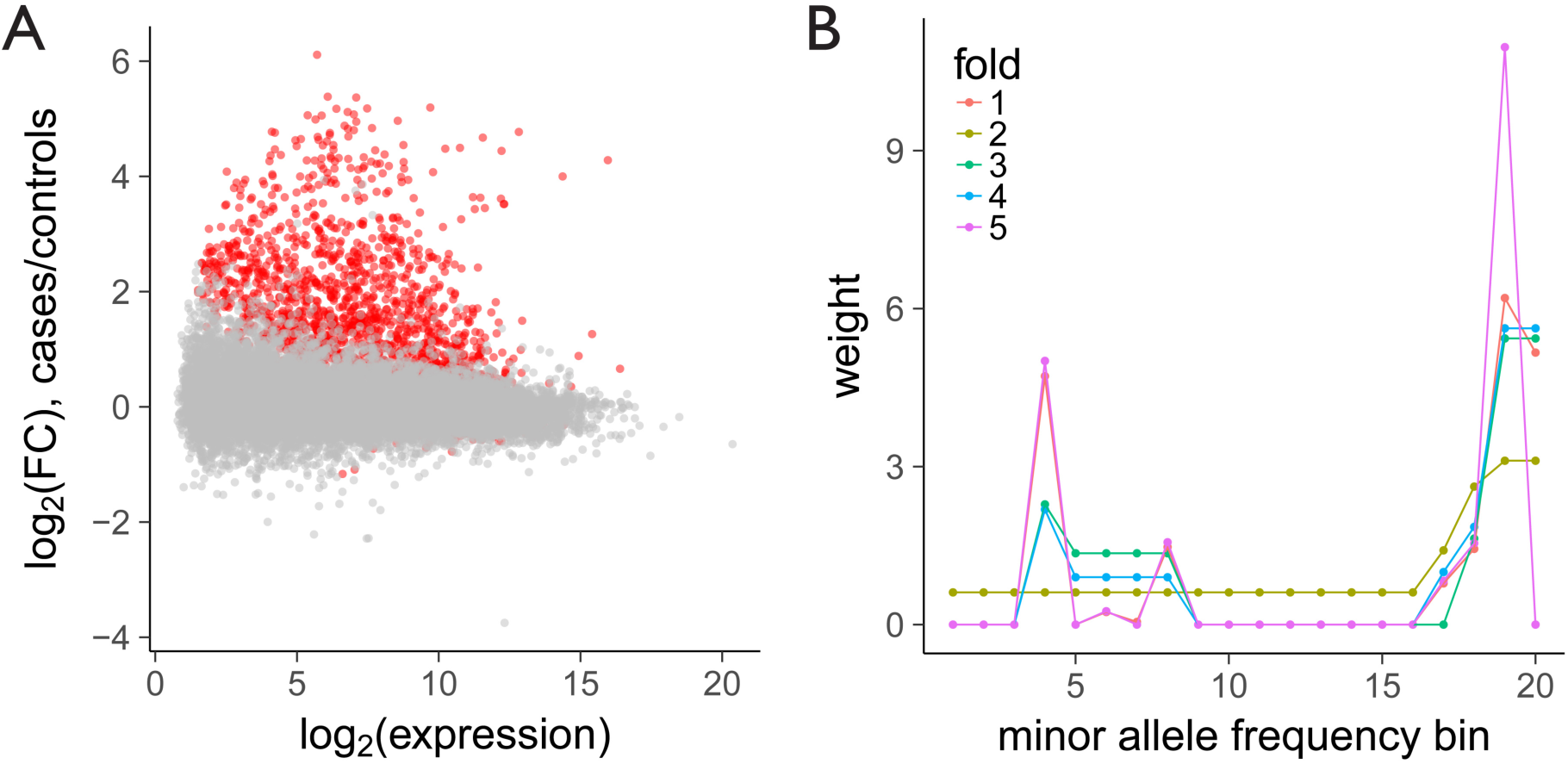
(**A**) Gene expression vs fold-change (FC) between GTEx Hashimoto Thyroiditis cases and controls. Genes with an adjusted p-value of less than 10^−7^ are colored red. (**B**) Independent Hypothesis Weighting (IHW) weights as a function of minor allele frequency bin. Each bin corresponds to a change of 0.01 in minor allele frequency between 0.1 and 0.5. Similar profiles for each fold demonstrate the robustness of the IHW fitting procedure.

**Figure S5.**
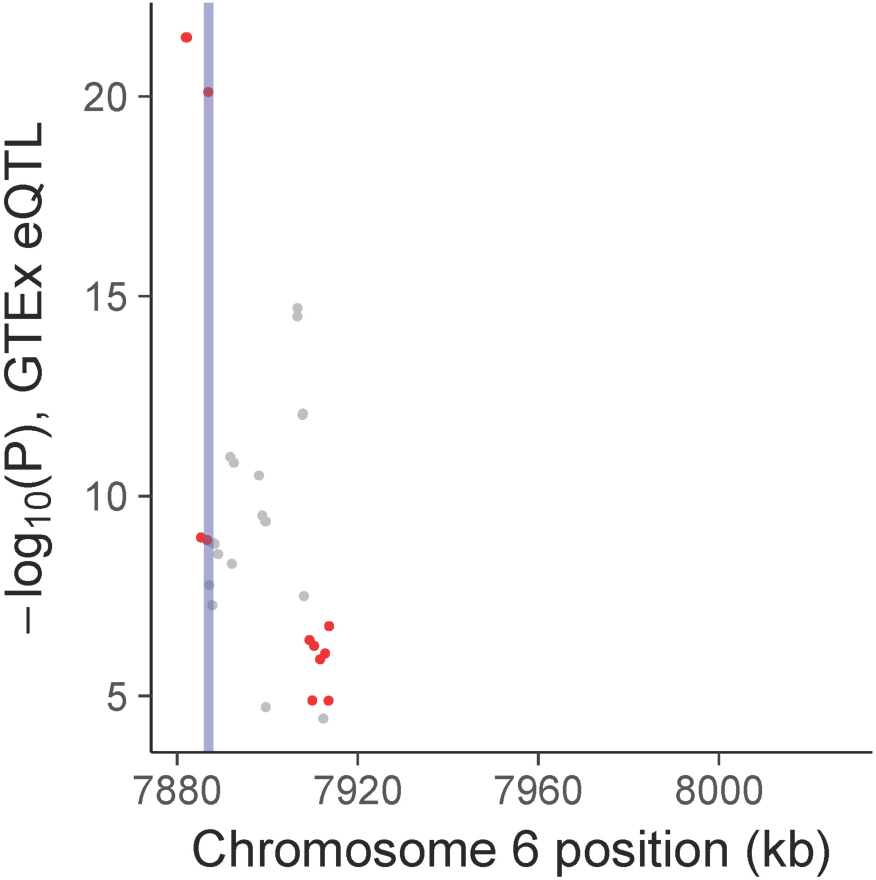
Significance of GTEx consortium v6p *cis*-expression QTLs for TXNDC5. Vertical blue line indicates the position of the top iQTL variant, *rs11962800*. Red points are eQTLs that are also iQTLs, or in linkage disequilibrium with iQTLs, as defined by a significance level of P_IHW_<0.4 (see Table S2). All significant eQTLs that were also tested in our iQTL analysis are shown. Genomic range is the same as in Figure 3D for comparison.

**Figure S6.**
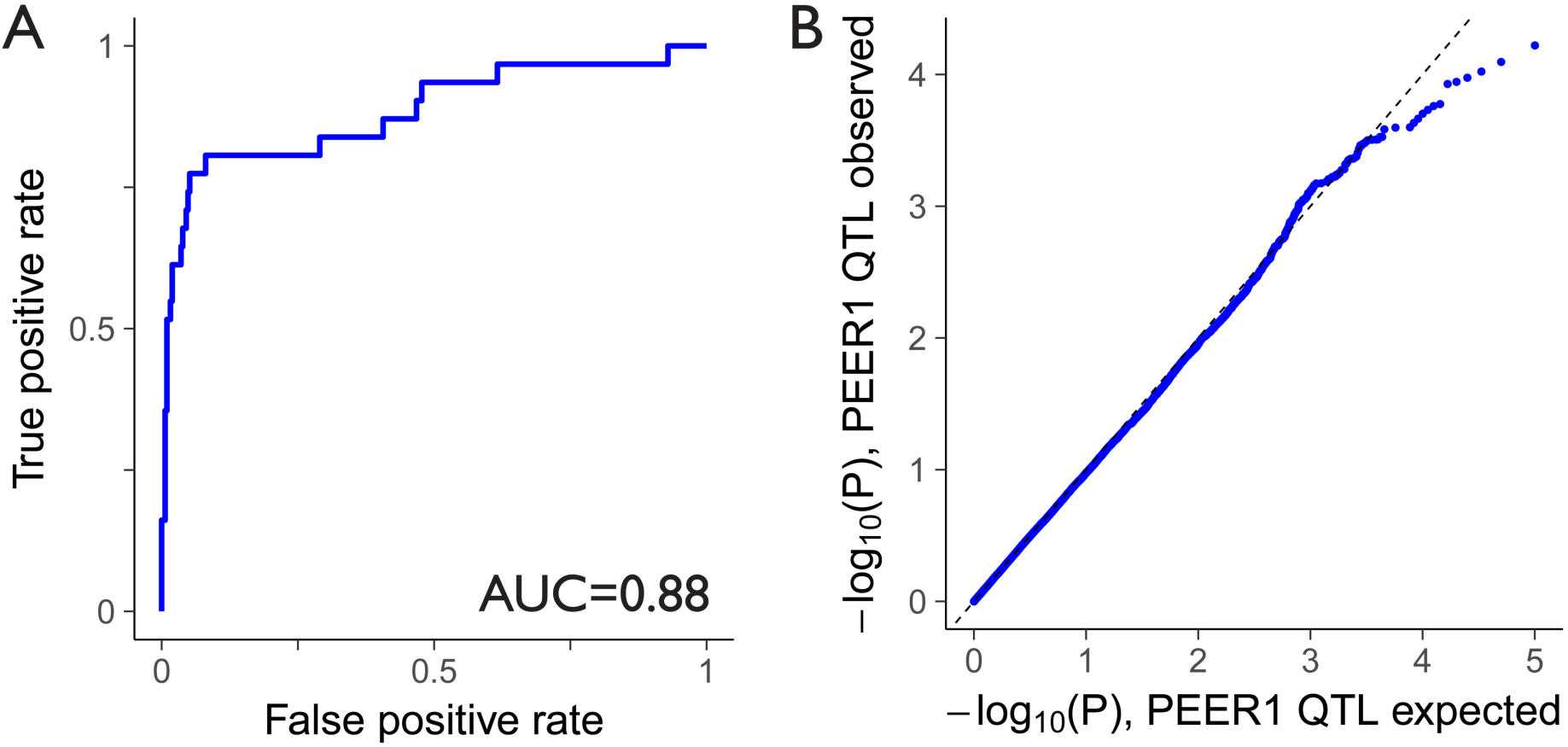
(**A**) ROC showing biomarker performance of the first expression PEER factor (PEER1) for predicting Hashimoto Thyroiditis. (**B**) A QQ plot showing expected vs observed p-values from PEER1 QTL fits of 100,215 candidate SNPs residing in the selected genes highlighted blue in Figure 3A.

**Figure S7.**
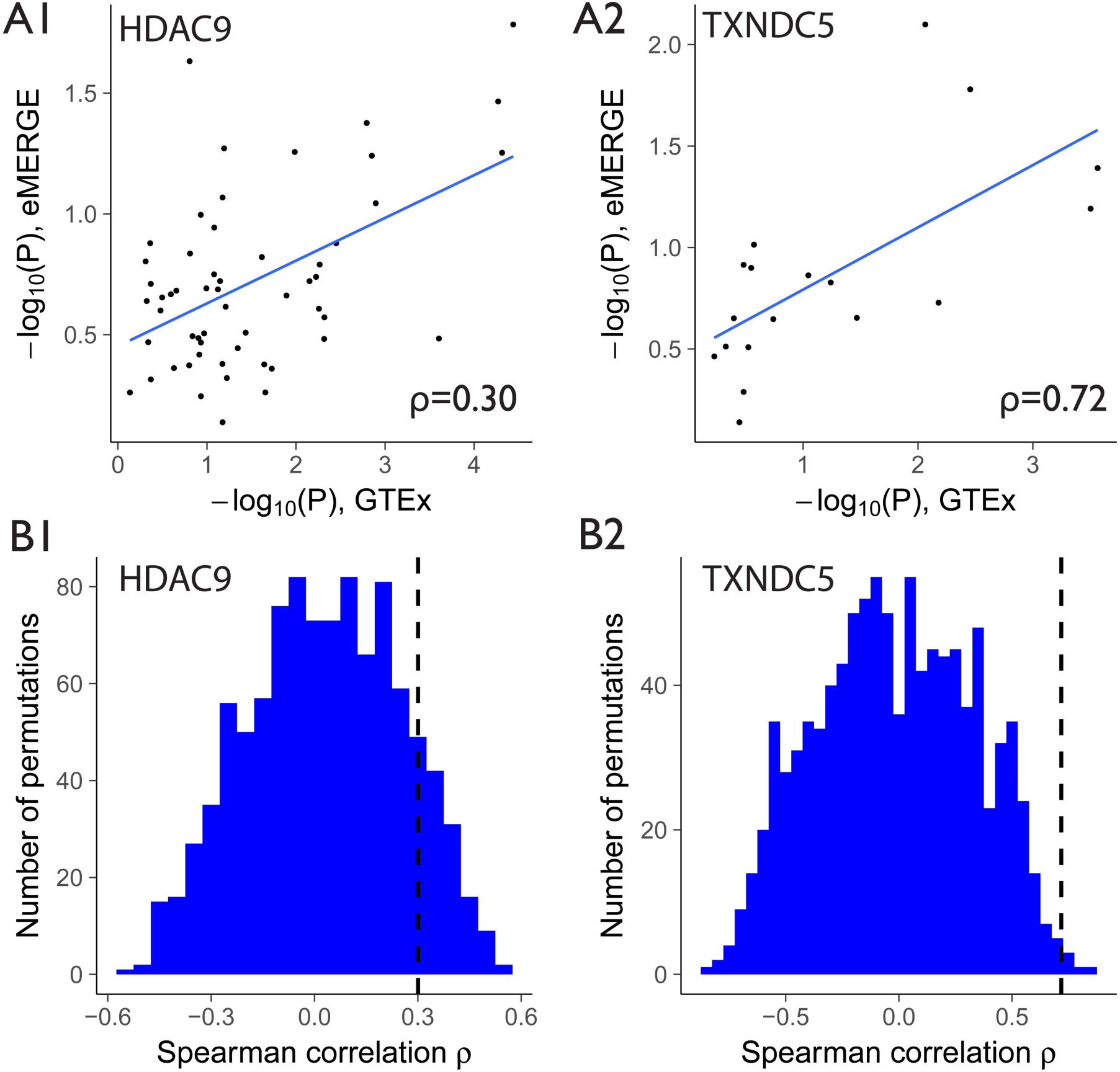
(**A1-2**) Concordance of the association signature between the GTEx and eMERGE data for HDAC9 (A1) and TXNDC5 (A2) for LD-pruned data. Each point represents the minimum p-value in each bin of size 15 SNPs for each dataset. (**B1-2**) Histograms of GTEx vs eMERGE correlation coefficients after 1,000 logistic regression analyses with permutations of the eMERGE hypothyroidism case/control labels. Dashed lines indicate Spearman correlation coefficients for the non-permuted data displayed in panels A1-2. Permutation p-values are 0.127 for HDAC9, and 0.006 for TXNDC5.

**Table S1.** Image QTLs with significant associations (P_IHW_<0.05) with our quantitative imaging biomarker for thyroid autoimmune disease.

| rsid | GRCh37/hg19 Coordinates | Gene | $P_{\text{IHW}}$ | FDR | P | $\beta$ | MAF | Haplotype |
| --- | --- | --- | --- | --- | --- | --- | --- | --- |
| rs17346782(C>T) | Chr7:18301455 | HDAC9 | 0.049 | 0.100 | 5.8E-06 | 2.50 | 0.17 | hap1 |
| rs12154277(G>T) | Chr7:18313167 | HDAC9 | 0.049 | 0.100 | 6.3E-06 | 1.91 | 0.43 | hap2 |
| rs12672535(G>A) | Chr7:18308913 | HDAC9 | 0.049 | 0.100 | 6.6E-06 | 1.90 | 0.44 | hap1 |
| rs6972958(G>T) | Chr7:18313358 | HDAC9 | 0.049 | 0.100 | 7.3E-06 | 1.90 | 0.44 | hap2 |
| rs6973137(G>A) | Chr7:18313517 | HDAC9 | 0.049 | 0.100 | 7.3E-06 | 1.90 | 0.44 | hap2 |
| rs4416737(C>T) | Chr7:18305489 | HDAC9 | 0.049 | 0.100 | 8.1E-06 | 1.91 | 0.43 | hap1 |
| rs73066383(C>A) | Chr7:18303874 | HDAC9 | 0.049 | 0.100 | 9.3E-06 | 1.90 | 0.43 | hap1 |
| rs12154412(G>A) | Chr7:18304326 | HDAC9 | 0.049 | 0.100 | 9.3E-06 | 1.90 | 0.43 | hap1 |
| rs4361680(T>A) | Chr7:18306817 | HDAC9 | 0.049 | 0.100 | 9.3E-06 | 1.90 | 0.43 | hap1 |
| rs7459237(C>A) | Chr7:18307448 | HDAC9 | 0.049 | 0.100 | 1.1E-05 | 1.89 | 0.43 | hap1 |
| rs302193(G>A) | Chr7:18298184 | HDAC9 | 0.049 | 0.100 | 1.1E-05 | 1.82 | 0.45 | hap1 |
| rs1107693(G>A) | Chr7:18307906 | HDAC9 | 0.049 | 0.100 | 1.2E-05 | 1.89 | 0.43 | hap1 |
| rs17138868(A>T) | Chr7:18294050 | HDAC9 | 0.049 | 0.113 | 1.5E-05 | 1.83 | 0.43 | hap1 |
| rs302178(G>A) | Chr7:18310791 | HDAC9 | 0.049 | 0.113 | 1.7E-05 | 1.80 | 0.44 | hap1 |
| rs73066378(G>A) | Chr7:18302581 | HDAC9 | 0.049 | 0.113 | 1.8E-05 | 1.89 | 0.42 | hap1 |
| rs7794278(G>A) | Chr7:18316661 | HDAC9 | 0.049 | 0.113 | 1.9E-05 | 1.82 | 0.43 | hap2 |
| rs17138874(T>C) | Chr7:18297439 | HDAC9 | 0.049 | 0.140 | 2.5E-05 | 1.79 | 0.46 | hap1 |
| rs6953847(C>T) | Chr7:18300086 | HDAC9 | 0.049 | 0.140 | 2.7E-05 | 1.80 | 0.44 | hap1 |
| rs11962800(A>G) | Chr6:7886905 | TXNDC5 | 0.049 | 0.166 | 4.5E-05 | -2.48 | 0.15 | hap3 |
| rs7779273(T>C) | Chr7:18301410 | HDAC9 | 0.049 | 0.210 | 7.4E-05 | 1.69 | 0.46 | hap1 |
| rs4721704(G>T) | Chr7:18301777 | HDAC9 | 0.049 | 0.262 | 1.1E-04 | 1.67 | 0.45 | hap1 |

**Table S2.**
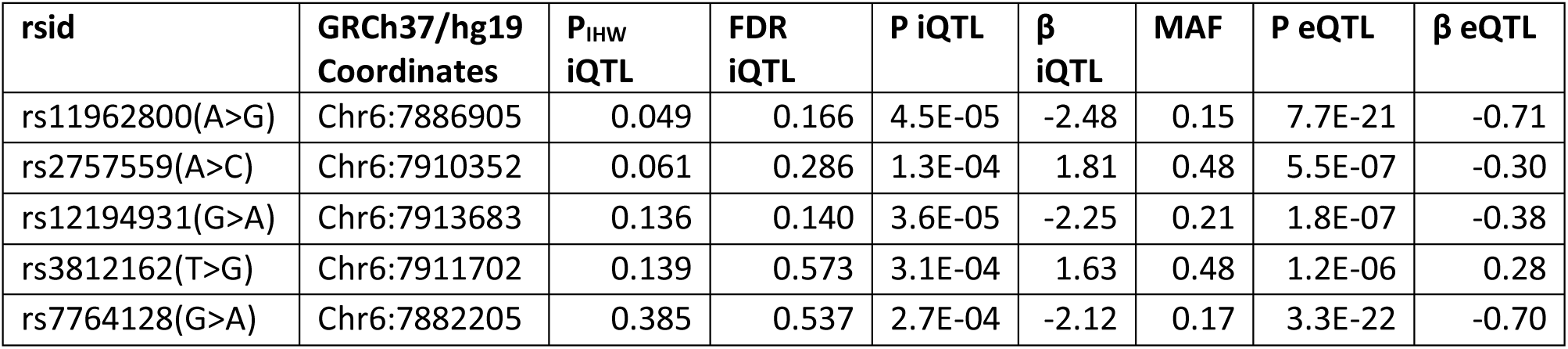
All iQTLs with P_IHW_<0.4 in TXNDC5 are also thyroid eQTLs in GTEx consortium data.

| rsid | GRCh37/hg19 Coordinates | $P_{\text{IHW}}$ iQTL | FDR iQTL | P iQTL | $\beta$ iQTL | MAF | P eQTL | $\beta$ eQTL |
| --- | --- | --- | --- | --- | --- | --- | --- | --- |
| rs11962800(A>G) | Chr6:7886905 | 0.049 | 0.166 | 4.5E-05 | -2.48 | 0.15 | 7.7E-21 | -0.71 |
| rs2757559(A>C) | Chr6:7910352 | 0.061 | 0.286 | 1.3E-04 | 1.81 | 0.48 | 5.5E-07 | -0.30 |
| rs12194931(G>A) | Chr6:7913683 | 0.136 | 0.140 | 3.6E-05 | -2.25 | 0.21 | 1.8E-07 | -0.38 |
| rs3812162(T>G) | Chr6:7911702 | 0.139 | 0.573 | 3.1E-04 | 1.63 | 0.48 | 1.2E-06 | 0.28 |
| rs7764128(G>A) | Chr6:7882205 | 0.385 | 0.537 | 2.7E-04 | -2.12 | 0.17 | 3.3E-22 | -0.70 |

## Transparent Methods

### Image Processing

341 publicly available GTEx histopathological thyroid images were downloaded from the Biospecimen Research Database (http://brd.nci.nih.gov/image-search/searchhome). For the purpose of segmenting individual tissue pieces only, the average intensity across color channels was calculated, and adaptive thresholding was performed to distinguish tissue from background. Interior holes in the tissue piece masks were filled using the *fillHull* function from the Bioconductor package *EBImage* [19]. In preparation for feature extraction, Gaussian blurring was used on each color channel to smooth out pixel-level variation on a length scale smaller than the observed lymphocyte invasion phenotypes. 117 textural Haralick features were then extracted using *EBImage* by calculating 13 base Haralick features for each of the three RGB color channels, and across three Haralick scales by sampling every 1, 10, or 100 pixels. Image metadata was extracted to verify identical pixel dimensions and scanning instrumentation across the dataset.

### Image Data Analysis

702 thyroid pieces were identified from the image segmentation. 49 pieces with overly small areas were removed. Averaged Haralick feature values were then calculated across tissue pieces for each sample. In preparation for downstream model fitting, each distribution was log_2_ transformed and rescaled using a Z-score. Five NA feature values were set to zero to avoid technical issues with the PCA fit.

### Gene Expression Analysis

GTEx v6 RNA-Seq data was downloaded from dbGaP (phs000424.v6.p1, 2015-10-05 release) under approved protocol #9112. Lowly expressed genes were defined as those which had a read count of less than 1 CPM in more than half of the samples, and were removed. Counts were log_2_ transformed after adding 1 to the counts to avoid the problem of taking the logarithm of zero. Quantile normalization was performed using the R package *limma*. Prior to running the gene expression data through CIBERSORT, RNA extraction batch was corrected for using *limma*. CIBERSORT analysis was performed at cibersort.stanford.edu after uploading gene expression data for the 341 GTEx thyroid samples. CIBERSORT default settings were used with 1,000 permutations and quantile normalization turned off since this step had been performed upstream. LM22 was chosen as the gene signature matrix. PEER factors were calculated using the R package *peer* (github: PMBio/peer). The fitting of image PCs against gene expression was performed using linear models in R with the formula *PC* = α + β*gene* + Σ_*k*_γ_k_*covariate*_*k*_ + ϵwhere covariates included the known confounders and PEER factors, as discussed in the main text. For each gene, p-values were extracted from *t*-statistics associated with the gene expression coefficient β. Differential expression analysis using *DESeq2* [26] was performed using the raw count data, with RNA extraction batch included as a technical covariate. As biological replicates were not available across samples, *DESeq2* estimated dispersion by pooling all samples.

### Genotype Analysis

GTEx genotype data was downloaded from dbGaP (phs000424.v6.p1, 2015-10-05 release) under approved protocol #9112. A total of 292 genotyped samples overlapped with the thyroid imaging samples. Minor allele frequency and missingness frequency filtering was performed using PLINK. QTL fits were performed using *MatrixEQTL*, which used the following model: *PC*2 = α + *βgenotype* + Σ_*k*_γ_k_*covariate*_*k*_ + ϵ. Genotype could take on the values 0/1/2/NA, and covariates were as described in the main text. QTL significance was extracted from the *t*-statistics associated with the genotype coefficient β. The correction for ancestry was performed by including as covariates the first 3 PCs of a PCA on the genotype matrix computed from the full set of 450 GTEx patients. Haplotypes were identified using PLINK, with a maximum block size of 5,000 kb. The haplotype identification utilized the full set of 450 GTEx samples. LD-pruning was performed using PLINK with a window size of 50 SNPs and an r^2^ threshold of 0.8. Reported genomic coordinates are for human genome build GRCh37/hg19.

eMERGE data was obtained from the study “eMERGE Network Imputed GWAS for 41 Phenotypes” (dbGaP study accession phs000888.v1.p1) under approved protocol #13896 for the consent cohorts Health/Medical/Biomedical (N=18,621) and Health/Medical/Biomedical-Genetic Studies Only-No Insurance Companies (N=15,911). Chromosome 6 and 7 genotype data from both cohorts was merged and 5,718 samples were selected based on their status as Chronic Autoimmune Hypothyroidism (accession phd004989.1) cases or controls. Inclusion criteria for hypothyroidism cases (C99269) included, but were not limited to, abnormal TSH/FT4 levels and the use of thyroid replacement medication. Exclusion criteria for hypothyroidism cases included secondary causes of hypothyroidism, hypothyroidism induced by surgery or radiation treatment, evidence of other thyroid diseases, or the use of thyroid-altering medication. Inclusion criteria for hypothyroidism controls (C99270) included no billing codes for hypothyroidism, no use of thyroid replacement medication, and normal TSH/FT4 levels. Exclusion criteria for controls included any evidence of hypo- or hyper-thyroidism, as well as other thyroid diseases, or the use of thyroid-altering medication. The Caucasian cohort was selected for further analysis, as described in the main text. Tests for Hardy-Weinberg Equilibrium (HWE) were performed using PLINK on the Caucasian cohort. One SNP in HDAC9 (*rs2058074*) had a HWE p-value less than 10^−6^ and was removed. Logistic regression models were fitted with the *glm* function in R using the following model: *logit*(*Y*) = α + *βgenotype* + Σ _*k*_γ_*k*_ *covariate*._*k*_ *Y* indicated the binary case/control disease status, genotype could take on the values 0/1/2/NA, and covariates were as described in the main text. The correction for ancestry was performed by including as covariates the first 3 PCs of a PCA on the genotype matrix computed from Chromosome 6 (for TXNDC9) and 7 (for HDAC9). Significance was extracted from the *z*-statistics associated with the genotype coefficient β. Colocalisation analysis was performed using the *coloc.abf* function from the R package *coloc*. Reported genomic coordinates are for human genome build GRCh37/hg19.

### Statistical Analysis

To test for associations between HT status and image PC2, we used the following logistic regression model: *logit*(*Y*) = α + β_1_*PC*2 + _2_ *sex* +:β _3_=*age* where *Y* indicated samples with HT phenotype according to GTEx pathology notes. The significance of association was extracted from the *z*-statistics associated with the beta coefficients. Due to co-linearity between sex and image PC2, associations between HT phenotype and sex were assessed using a reduced model without the image PC2 term. Enrichment of fibrotic phenotypes within the set of samples with HT phenotypes was assessed using Fisher’s Exact Test. Gene-set enrichment tests were performed using a Fisher’s Exact Test for each of the 13,776 tested Gene Ontology terms. Multiple-testing correction across terms was performed using the method of Benjamini and Hochberg. Independent Weighting Hypothesis p-value correction on the image QTL results was performed using the Bioconductor software *IHW* [28]. The permutation p-values reported for the eMERGE validations were estimated using the formula (*M* + 1)(+ 1) where *M* is the number of permutations with a correlation coefficient exceeding that of the non-permuted data, and *N* is the total number of permutations.

## Reference

1. Pickrell, J.K., et al., Understanding mechanisms underlying human gene expression variation with RNA sequencing. Nature, 2010. 464: p. 768–772.

2. Westra, H.-J., et al., Systematic identification of trans eQTLs as putative drivers of known disease associations. Nature Genetics, 2013. 45: p. 1238–1243.

3. Bulik-Sullivan, B., et al., An atlas of genetic correlations across human diseases and traits. Nature Genetics, 2015. 47: p. 1236–1241.

4. Gutierrez-Arcelus, M., S.S. Rich, and S. Raychaudhuri, Autoimmune diseases — connecting risk alleles with molecular traits of the immune system. Nature Reviews Genetics, 2016. 17: p. 160–174.

5. Chalasani, N., et al., Genome-Wide Association Study Identifies Variants Associated with Histologic Features of Nonalcoholic Fatty Liver Disease. Gastroenterology, 2010. 139: p. 1567–1576.e6.

6. Layfield, L.J., et al., Accuracy and Reproducibility of Nuclear/Cytoplasmic Ratio Assessments in Urinary Cytology Specimens. Diagnostic Cytopathology, 2017. 45: p. 107–112.

7. Qureshi, A., et al., Gleason’s Grading of Prostatic Adenocarcinoma: Inter-Observer Variation Among Seven Pathologists at a Tertiary Care Center in Oman. Asian Pacific journal of cancer prevention: APJCP, 2016. 17: p. 4867–4868.

8. Colquhoun, P., et al., Interobserver and intraobserver bias exists in the interpretation of anal dysplasia. Diseases of the Colon and Rectum, 2003. 46: p. 1332–1336; discussion 1336-1338.

9. Yu, K.-H., et al., Predicting non-small cell lung cancer prognosis by fully automated microscopic pathology image features. Nature Communications, 2016. 7.

10. Khan, F.M., et al., Predicting and replacing the pathological Gleason grade with automated gland ring morphometric features from immunofluorescent prostate cancer images. Journal of Medical Imaging (Bellingham, Wash.), 2017. 4: p. 021103.

11. Cho, M.H., et al., A Genome-Wide Association Study of Emphysema and Airway Quantitative Imaging Phenotypes. American Journal of Respiratory and Critical Care Medicine, 2015. 192: p. 559–569.

12. Shen, L., et al., Whole genome association study of brain-wide imaging phenotypes for identifying quantitative trait loci in MCI and AD: A study of the ADNI cohort. NeuroImage, 2010. 53: p. 1051–1063.

13. Popovici, V., et al., Joint analysis of histopathology image features and gene expression in breast cancer. BMC Bioinformatics, 2016. 17: p. 209.

14. Yuan, Y., et al., Quantitative Image Analysis of Cellular Heterogeneity in Breast Tumors Complements Genomic Profiling. Science Translational Medicine, 2012. 4: p. 157ra143–157ra143.

15. Beck, A.H., et al., Systematic Analysis of Breast Cancer Morphology Uncovers Stromal Features Associated with Survival. Science Translational Medicine, 2011. 3: p. 108ra113–108ra113.

16. Consortium, G., Human genomics. The Genotype-Tissue Expression (GTEx) pilot analysis: multitissue gene regulation in humans. Science (New York, N.Y.), 2015. 348: p. 648–660.

17. Lonsdale, J., et al., The Genotype-Tissue Expression (GTEx) project. Nature Genetics, 2013. 45: p. 580– 585.

18. Pyzik, A., et al., Immune disorders in Hashimoto’s thyroiditis: what do we know so far? Journal of Immunology Research, 2015. 2015: p. 979167.

19. Pau, G., et al., EBImage—an R package for image processing with applications to cellular phenotypes. Bioinformatics, 2010. 26: p. 979–981.

20. Haralick, R.M., Statistical and structural approaches to texture. Proceedings of the IEEE, 1979. 67: p. 786– 804.

21. Anila, K., N. Nayak, and K. Jayasree, Cytomorphologic spectrum of lymphocytic thyroiditis and correlation between cytological grading and biochemical parameters. Journal of Cytology / Indian Academy of Cytologists, 2016. 33: p. 145–149.

22. Iannaci, G., et al., Fibrous Variant of Hashimoto’s Thyroiditis as a Diagnostic Pitfall in Thyroid Pathology. Case Rep Endocrinol, 2013. 2013: p. 308908.

23. Stegle, O., et al., Using probabilistic estimation of expression residuals (PEER) to obtain increased power and interpretability of gene expression analyses. Nature Protocols, 2012. 7: p. 500–507.

24. Aguet, F., et al., Local genetic effects on gene expression across 44 human tissues. bioRxiv, 2016: p. 074450.

25. Newman, A.M., et al., Robust enumeration of cell subsets from tissue expression profiles. Nature Methods, 2015. 12: p. 453–457.

26. Love, M.I., W. Huber, and S. Anders, Moderated estimation of fold change and dispersion for RNA-seq data with DESeq2. Genome Biology, 2014. 15: p. 550.

27. Shabalin, A.A., Matrix eQTL: ultra fast eQTL analysis via large matrix operations. Bioinformatics (Oxford, England), 2012. 28: p. 1353–1358.

28. Ignatiadis, N., et al., Data-driven hypothesis weighting increases detection power in genome-scale multiple testing. Nature Methods, 2016. 13: p. 577–580.

29. Yan, K., et al., Histone Deacetylase 9 Deficiency Protects against Effector T Cell-mediated Systemic Autoimmunity.The Journal of Biological Chemistry, 2011. 286: p. 28833–28843.

30. Kim, D.W., et al., A histone deacetylase inhibitor improves hypothyroidism caused by a TRα1 mutant. Human Molecular Genetics, 2014. 23: p. 2651–2664.

31. Tang, J., H. Yan, and S. Zhuang, Histone deacetylases as targets for treatment of multiple diseases. Clinical Science, 2013. 124: p. 651–662.

32. Beck, T., et al., GWAS Central: a comprehensive resource for the comparison and interrogation of genome-wide association studies. European journal of human genetics: EJHG, 2014. 22: p. 949–952.

33. Anderson, C.A., et al., Meta-analysis identifies 29 additional ulcerative colitis risk loci, increasing the number of confirmed associations to 47. Nature Genetics, 2011. 43: p. 246–252.

34. Shizuma, T., Concomitant Thyroid Disorders and Inflammatory Bowel Disease: A Literature Review. BioMed Research International, 2016. 2016: p. e5187061.

35. Sullivan, D.C., et al., EndoPDI, a novel protein-disulfide isomerase-like protein that is preferentially expressed in endothelial cells acts as a stress survival factor. The Journal of Biological Chemistry, 2003. 278: p. 47079–47088.

36. Chang, X., et al., Investigating a pathogenic role for TXNDC5 in rheumatoid arthritis. Arthritis Research & Therapy, 2011. 13: p. R124.

37. Jeong, K.-H., et al., Association of TXNDC5 gene polymorphisms and susceptibility to nonsegmental vitiligo in the Korean population. British Journal of Dermatology, 2010. 162: p. 759–764.

38. McCall, Matthew N., Peter B. Illei, and Marc K. Halushka, Complex Sources of Variation in Tissue Expression Data: Analysis of the GTEx Lung Transcriptome. The American Journal of Human Genetics, 2016. 99: p. 624–635.

39. Gottesman, O., et al., The Electronic Medical Records and Genomics (eMERGE) Network: past, present, and future. Genetics in Medicine, 2013. 15: p. 761–771.

40. Denny, Joshua C., et al., Variants Near FOXE1 Are Associated with Hypothyroidism and Other Thyroid Conditions: Using Electronic Medical Records for Genome- and Phenome-wide Studies. The American Journal of Human Genetics, 2011. 89: p. 529–542.

